# Bottom-up ecology of the human microbiome: from metagenomes to metabolomes

**DOI:** 10.1101/060673

**Authors:** Daniel R. Garza, Marcel C. Van Verk, Martijn A. Huynen, Bas E. Dutilh

## Abstract

The environmental metabolome is a dominant and essential factor shaping microbial communities. Thus, we hypothesized that metagenomic datasets could reveal the quantitative metabolic status of a given sample. Using a newly developed bottom-up ecology algorithm, we predicted high-resolution metabolomes of hundreds of metagenomic datasets from the human microbiome, revealing body-site specific metabolomes consistent with known metabolomics data, and suggesting that common cosmetics ingredients are some of the major metabolites shaping the human skin microbiome.

## Main text

Microbial communities constantly adapt to exploit available resources [Martiny 2015]. As a result, the presence of specific microbes or distributions of microbes allows us to infer environmental features. For example, the altered metabolic conditions in the microenvironment of colorectal cancer (CRC) tumors select for the outgrowth of specific species in the human CRC microbiome [Tjalsma 2012], allowing cancer detection [Zeller 2014]. Similarly, microbes can serve as biosensors for geochemical features such as solvent or uranium contamination [Smith 2015]. These and many other empirical examples of significant associations between the environment and the microbiota [Merrifield 2016, Adams 2015] suggest that the composition and metabolic potential of microbial communities could be used to reconstruct the metabolic environment through a reverse engineering strategy.

Shotgun metagenomics rapidly inventories the composition and genomic content of microbiomes, but obtaining similarly high-resolution metabolomic measurements of microbial environments remains challenging, among others due to the hundreds to thousands of biochemical compounds that may be difficult to distinguish using commonly used mass spectrometry-based technologies [Marcobal 2013]. Here we address this challenge by developing a novel approach that predicts the metabolic environment of a microbial community based on its metagenome, by using the inferred metabolism and abundance profiles of community members as input.

The main premise of our approach, which we named bottom-up ecology, is that the metabolic potential of a microbiome that is encoded in the genes of the species and their wiring into pathways, reveals how it can exploit the available metabolic resources. Using bottom-up ecology, we can quantitatively infer these resources by identifying the metabolic environment that yields growth of these microbes in the relative abundances as observed in the metagenome. Our approach is outlined in Figure 1 (see Supplementary Methods for details). First, we use reference genome sequences to generate genome-scale metabolic models of the species encountered (GSMMs) by using an established automated pipeline (1) [Henry 2010]. GSMMs provide a minimally biased description of the metabolic preferences of a microbial genome by integrating prior knowledge about protein functions based on information from many model and non-model organisms, and can be used to model the fluxes through the cellular metabolic network [Henry 2010]. Next, we determine the relative abundances of these organisms in the microbiome by mapping the metagenomic reads against the reference genomes (2). We can now ask the question, which metabolic environment would lead to relative growth rates of the GSMMs (3) that best correlate with the observed relative abundances (4), where growth is defined as the sum of all fluxes in biomass reactions in the GSMMs [Henry 2010]. Flux balance analysis (FBA) allows the growth or biomass production of GSMMs to be estimated given certain constraints. We constrain the GSMMs of all the microbes found in a metagenome by providing the same metabolic environment (modeled as an upper bound to the import reactions of metabolites) and assuming the same cellular objective of growth. Finally, we use semi-Markov chain sampling of the highly dimensional metabolome solution space to identify the metabolomic composition leading to the GSMM growth profile that optimally correlates with the abundance profile of the microbial genomes observed in the metagenome (5).

**Figure 1.**
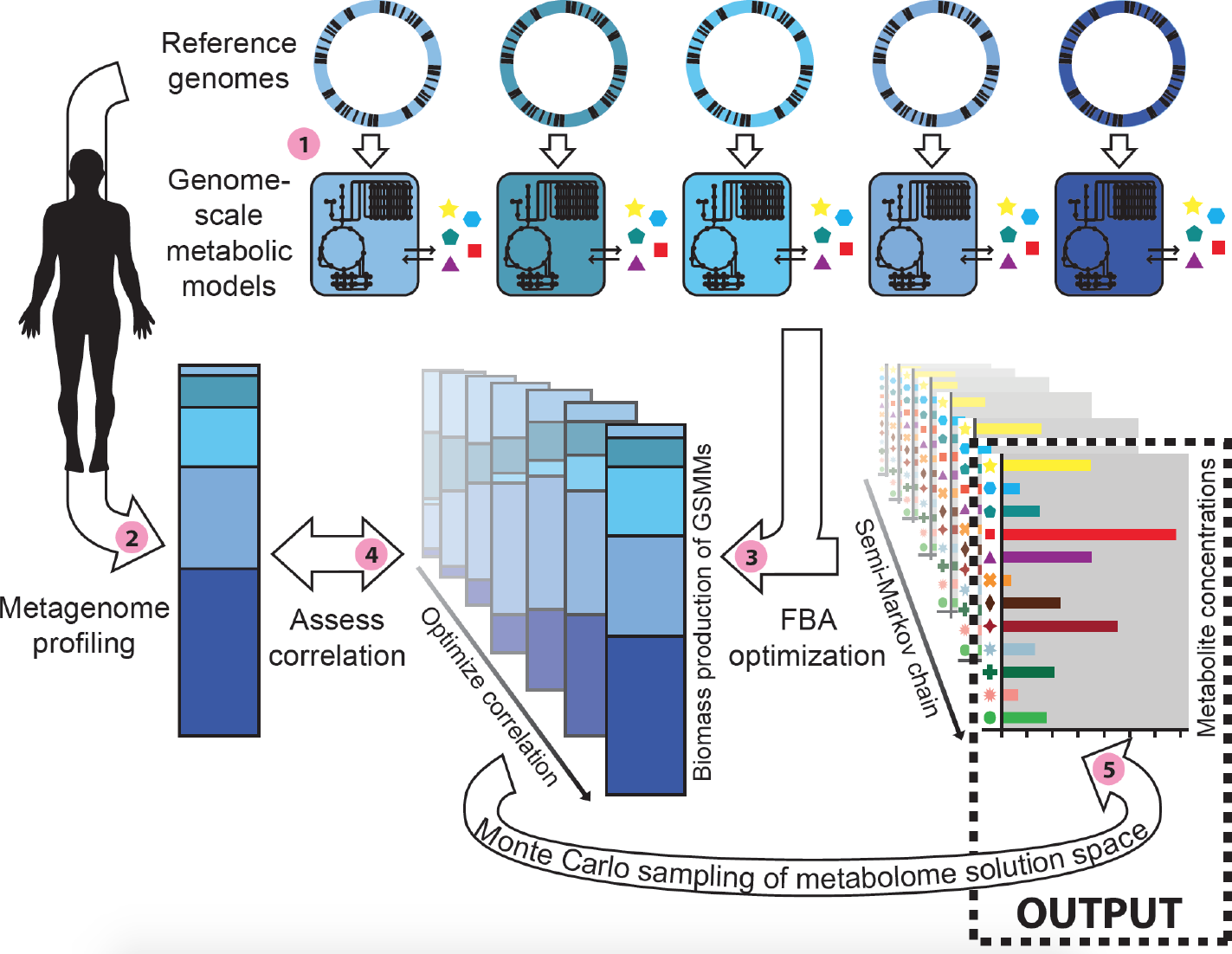
*Algorithm for bottom-up ecology from metagenomes to metabolomes. See text for details.*

To test our bottom-up ecology approach in a well-controlled system, we generated nine *in silico* metabolic environments by sampling random metabolites according to a realistic distribution (see Supplementary Methods). Next, we inferred growth of GSMMs generated from the bacterial reference genomes of the Human Microbiome Project (HMP) [Consortium 2012], and assumed that the microbes with the highest growth rates would be the most abundant in the simulated metagenomes. Using only the predicted abundance values of the best growing species, we then re-inferred the environment based on their relative abundance profiles. As expected, the metabolomes inferred by using bottom-up ecology correlate with the metabolomes on which the species were grown, while this correlation is absent when comparing metabolomes predicted from non-matching species (Figure 2a). Importantly, correct inference of the metabolic environment depends on the information from the entire simulated microbial community. For example, the correlation values on the diagonal in the top-20 species plot were significantly lower than in the top-100 species plot (P-value=0.04, one-tailed paired T-test of nine values, see Figure 2a).

**Figure 2.**
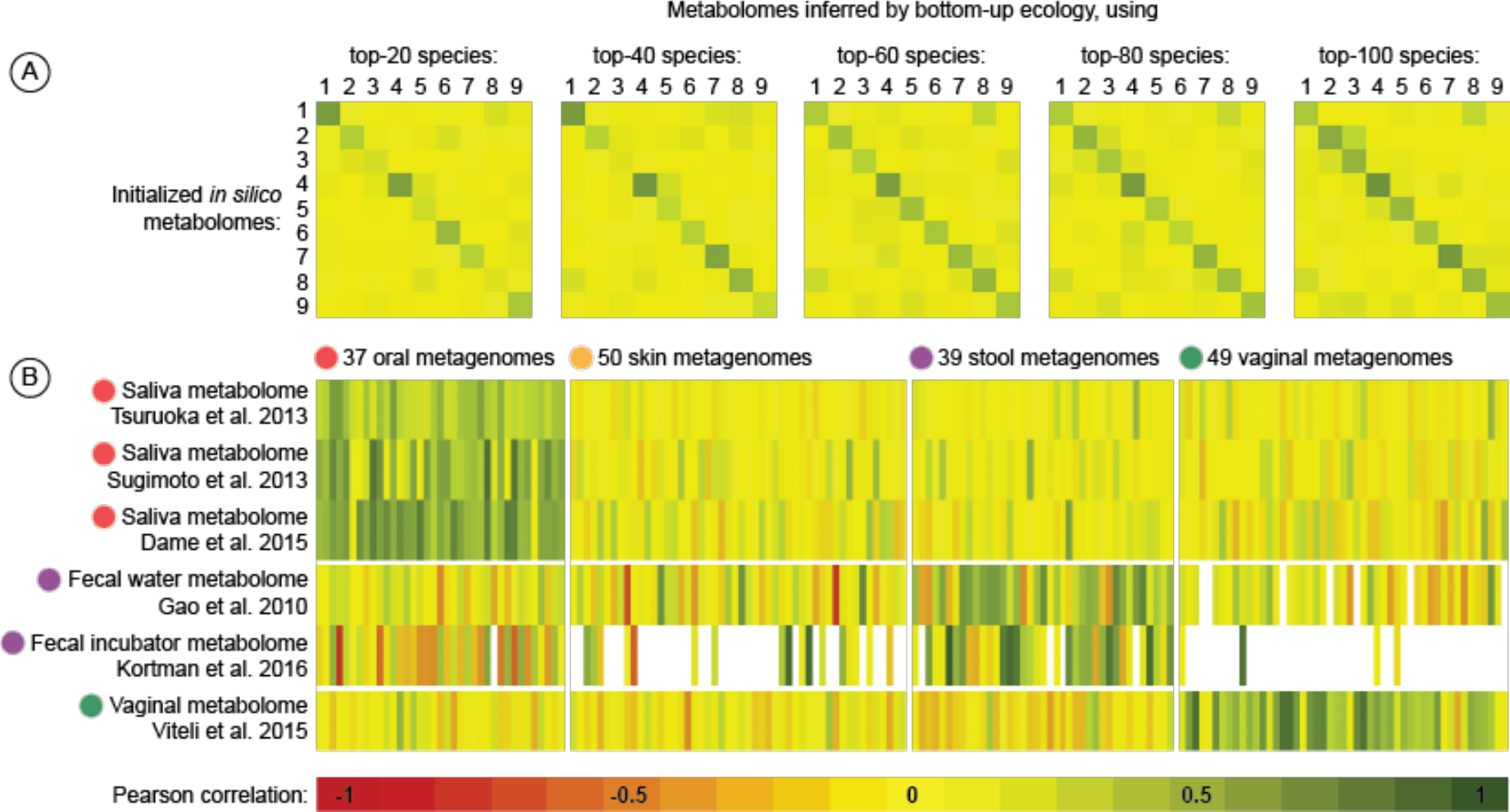
*Pearson correlations between true and predicted metabolomic profiles. In A, the “true” metabolomes consist of nine random metabolic environments created in silico, while the predicted metabolomes were inferred using the top-20, top-40, top-60, top-80, and top-100 best growing species in those environments. In B, the “true” metabolomes consist of six experimentally measured metabolomes from literature, while the predicted metabolomes were inferred using 175 HMP metagenomes from four body-sites. Correlations are only shown if >5 metabolites of the predicted metabolites were measured and vice versa. See Supplementary File 2 for details.*

Next, we tested the ability of the bottom-up ecology algorithm to infer the metabolomic environment in multiple human body-sites. We used 175 metagenomic datasets from four different body-sites (oral, skin, stool, and vaginal) where the relative abundances were measured of over one thousand reference bacteria [Consortium 2012]. A principal component analysis of the resulting predicted metabolomic environments revealed four clusters corresponding to the origin of the metagenomes (Supplementary Figure 1a), as was reflected in the body-site specific microbiomes that were previously observed [Consortium 2012]. This clustering was independent of the initialization of our algorithm in the highly dimensional metabolome search space. For example, searches based on oral metagenomes that were initiated with predicted skin metabolomes quickly converged to the oral metabolome cluster (Supplementary Figures 1b–e). Thus, our bottom-up ecology algorithm predicted consistent and specific metabolomes for the human microbiome body-sites.

The metabolomes inferred from the HMP metagenomes allowed us to identify the most important metabolites in the different human body-sites. Due to the difficulties in obtaining quantitative, high-resolution metabolomic profiles of a sample [Marcobal 2013], there are relatively few experimental datasets that could be used for experimental benchmarking of our bottom-up ecology algorithm. From the Human Metabolome Database [Wishart 2013], we obtained three annotated metabolomes from saliva and one from fecal water. Moreover, we included one additional metabolome from a fecal incubator and one from vagina from recent literature [Kortman 2016, Vitali 2015]. We linked measured metabolites between these studies and our models (Supplementary File 1) and correlated the metabolomic profiles of these six independently measured metabolomes to the values of the metabolites inferred by bottom-up ecology from the 175 HMP metagenomic datasets. As shown in Figure 2b, the oral metagenomes are linked to the saliva metabolomes, the stool metagenomes to the fecal water and fecal incubator metabolomes, and the vaginal metagenomes to the vaginal metabolome (P-value≈0, one-tailed unpaired T-test). These results show that the metabolomic environment of a microbiome can be quantitatively captured in high-resolution by our bottom-up ecology algorithm.

To our knowledge, no high-throughput human skin metabolome has been measured to date. To identify the main metabolites on the human skin, we averaged the relative concentrations of all metabolites in the metabolomes predicted based on 50 skin metagenomes (Supplementary File 1). Interestingly, the top compounds shaping the skin microbiome include various ingredients found in cosmetic and hygiene products (Supplementary Table 1). For example, myristic acid is used as a fragrance ingredient, cleansing agent, and emulsifier, and is readily adsorbed by the skin. Similarly, citrate is a commonly used ingredient to adjust the acidity of cosmetics. Nicotinamide ribonucleotide, aspartate, and N-acetyl glucosamine are used in skin conditioner products. Moreover, N-acetyl glucosamine is a precursor to hyaluronic acid, a major component of skin structure, a pathway that responds to UV irradiation in skin [Averbeck 2007]. These results provide a first look into the metabolites shaping the human skin microbiome [Bouslimani 2015, Grice 2011].

Our bottom-up ecology approach exploits the fact that in an ecosystem, microbes are constantly competing for resources, leading to a relative abundance distribution reflecting their ability to exploit these resources. Metagenome-guided modeling enables a deeper understanding of microbial ecosystems by linking the environmental metabolome to the metabolic network of individual microbial populations [Levy 2013]. By explicitly modeling the fluxes of individual GSMMs that are matched with the species composition of the system in a probabilistic fashion, our approach provides a starting point for mechanistic models of microbial ecology, including the potential for systems with more complex crossfeeding networks [Garza 2015].

While our results show meaningful metabolomic profiles for diverse human body-site environments, there is still a relatively high level of noise (Figure 2b). First, it should be noted that the metagenomes and metabolomes used in this benchmark were measured and published independently in systems that may differ in unknown ways, e.g. a fecal incubator versus fresh stool. Second, automatically reconstructed GSMMs have ~70% agreement with experimentally measured microbial phenotypes [Henry 2010, Plata 2015], adding significant noise to the inferred metabolomic profiles. Third, our algorithm depends on the availability of reference genome sequences to derive these GSMMs, as do approaches that exploit environmental sequencing data to make inferences about the genetic functionality of a microbial ecosystem [Langille 2013, Aßhauer 2015]. Ongoing work addressing these points is expected to improve bottom-up ecology as an approach to predict metabolic features of microbial environments, also in environments that are less well-sampled than the human microbiome. Nevertheless, it is promising that we can already reconstruct the metabolomes of human body-site environments by starting only from the metagenomic content of the microbial community.

## Supplementary Methods

### Datasets

From the US Department of Energy Systems Biology Knowledgebase (KBase, www.kbase.us) and the Human Microbiome Project (HMP, www.hmpdacc.org) [Consortium 2012, Martin 2012] we downloaded the human microbiome reference genomes and their abundance profiles in 175 metagenomes, respectively, including 37 oral, 50 skin, 39 stool, and 49 vaginal metagenomes (listed in Supplementary File 1). Experimentally measured metabolomic profiles were obtained from the Human Metabolomics Database [Wishart 2013], including one from fecal water [Gao 2010] and three from saliva [Tsuruoka 2013, Sugimoto 2013, Dame 2015]. Additionally, one stool [Kortman 2016] and one vaginal [Vitali 2015] metabolome were obtained from recent literature.

### Metabolic modeling

We used the ModelSEED pipeline [Henry 2010] to generate genome-scale metabolic models (GSMMs) for the 1,145 HMP reference genomes that were present in at least one of the metagenomic datasets. Briefly, genomic annotations were used to identify the biochemical reactions in a species’ metabolic network. The molecular stoichiometry of these reactions was expressed in a matrix that transforms reaction-rates to the time-derivative of metabolite concentrations. The nullspace of this matrix contains the equilibria solutions for reaction rates. Parsimonious gapfilling was applied by adding the minimal possible set of reactions to the model that are essential for a model to grow, i.e. to yield a flux through the biomass reactions [Henry 2010]. Gap-filled reactions were likely missed during sequencing, assembly, or genome annotation. We excluded dead-end exchange reactions from the models that remained unresolved after gap filling or had no influence on the objective function. Flux balance analysis (FBA) simulations were performed in a Python 2.7 environment, using the COBRApy package for constraint-based modeling [Ebrahim 2013] and Gurobi 5.6.3 (www.gurobi.com) or GLPK 4.35 (www.gnu.org/software/glpk) as linear programming solvers. To reflect the constant competition between microbes, we used growth as the objective function in the FBA [Orth 2010].

### Bottom-up ecology algorithm

The input of our bottom-up ecology approach (outlined in Figure 1 and main text) are (i) a list of microbes and their relative abundances, and (ii) a database of GSMMs generated from the genomes of these microbes. Thus, the approach depends on the availability of high-quality draft reference genome sequences, as are available for the microbes found in the human microbiome and increasingly also for other environments. Typically, one GSMM will have 35-80 exchange reactions representing the metabolic compounds that the organism can utilize. Depending on the complexity of the microbiome, the GSMMs of all the microbes in a community together will be able to utilize >200 different metabolites. These combined exchange reactions represent the metabolites whose environmental concentrations are inferred by the bottom-up ecology framework.

At the core of the approach is an optimization algorithm that searches the >200-dimensional metabolome search space for a composition of the metabolomic environment that, when applied simultaneously to the GSMMs of all co-existing microbes using FBA, yields a relative biomass production profile *b* that correlates with the abundance profile *m* of the microbes in the metagenome. The metabolic compound concentrations are modeled in the FBA as an upper bound to the influx reaction. We use Monte Carlo optimization following a semi-Markov chain to search the highly dimensional solution space. After random initialization (or initialization with a decoy metabolome as in Supplementary Figures 1b–e), a new candidate environment *e’* is generated from the current environment *e*, by slightly altering the concentration of one metabolite following a uniform distribution. The maximum biomass production rates of all GSMMs are then evaluated for the candidate environment, and the change is accepted if the Pearson correlation of the metagenomic abundances with the growth rates in the candidate environment, ρ(*m,b_e’_*) is higher than for the current environment ρ(*m,b_e_*), or with a uniform probability ρ(*m,b_e’_*)/ρ(*m,b_e_*) otherwise. Every 150 search steps, the algorithm evaluates the past outcomes and chooses the environment that yielded the highest correlation [Barbu 2008]. Samples are first subjected to 100,000 search steps, and 100,000 steps are subsequently added until a high Pearson correlation (ρ>0.6) with the target metagenomic abundance profile is achieved. Finally, the 10% time points with the highest Pearson correlation scores between the biomass profile and the metagenomic abundance profile are averaged and constitute the predicted metabolome. Note that the correlation that is optimized using the semi-Markov chain is the correlation ρ(m,b_e_) between the metagenomic species profile *m* and the biomass production rates *b_e_,* while the correlations that are shown in our results (e.g. in Figure 2b) are correlations between the predicted metabolome *e* and experimentally measured metabolic concentrations.

The Cython/Python implementation of the bottom-up ecology algorithm are available at https://github.com/danielriosgarza/BottomUpEcologyFunctions.

### In silico random metabolomes

To test the bottom-up ecology algorithm in a well-controlled system, we generated nine random metabolomes *in silico.* To implement a realistic metabolite distribution in these *in silico* metabolomes, we first used maximum likelihood estimates to assess the shape, location, and scale parameters that most accurately fit the distribution of metabolite abundances in all 175 predicted metabolomes. We fitted 74 common statistical distributions and found that most of the metabolomes could be modeled by right-skewed positive distributions, the Burr (type III) distribution resulting in the lowest average square mean errors (Supplementary Figure 2). Thus, nine random metabolic environments were drawn according to this distribution, and GSMMs based on all HMP reference genomes were grown on each *in silico* metabolome and sorted by growth rate. Finally, metabolomes were re-inferred based on the relative abundance profiles of the top-20, top-40, top-60, top-80, and top-100 best-growing species.

### Statistical analysis

Statistical analysis, including the maximum likelihood estimate for statistical distributions were performed on a Python 2.7 environment, using the “stat” statistical package of Scipy 0.15.1. Principal coordinate analyses were performed using the scikit-learn package.

## Author contributions

DRG created the algorithm and performed the experiments. All authors devised the study and wrote the manuscript.

## Acknowledgments

We thank Maarten Kooyman from SURFsara for help implementing bottom-up ecology on the Netherlands Life Science Grid. DRG is supported by the Science Without Borders program of CNPQ/BRASIL. BED is supported by Netherlands Organization for Scientific Research (NWO) Vidi grant 864.14.004.

## Supplementary Figures, Files, and Table

**Supplementary Figure 1.**
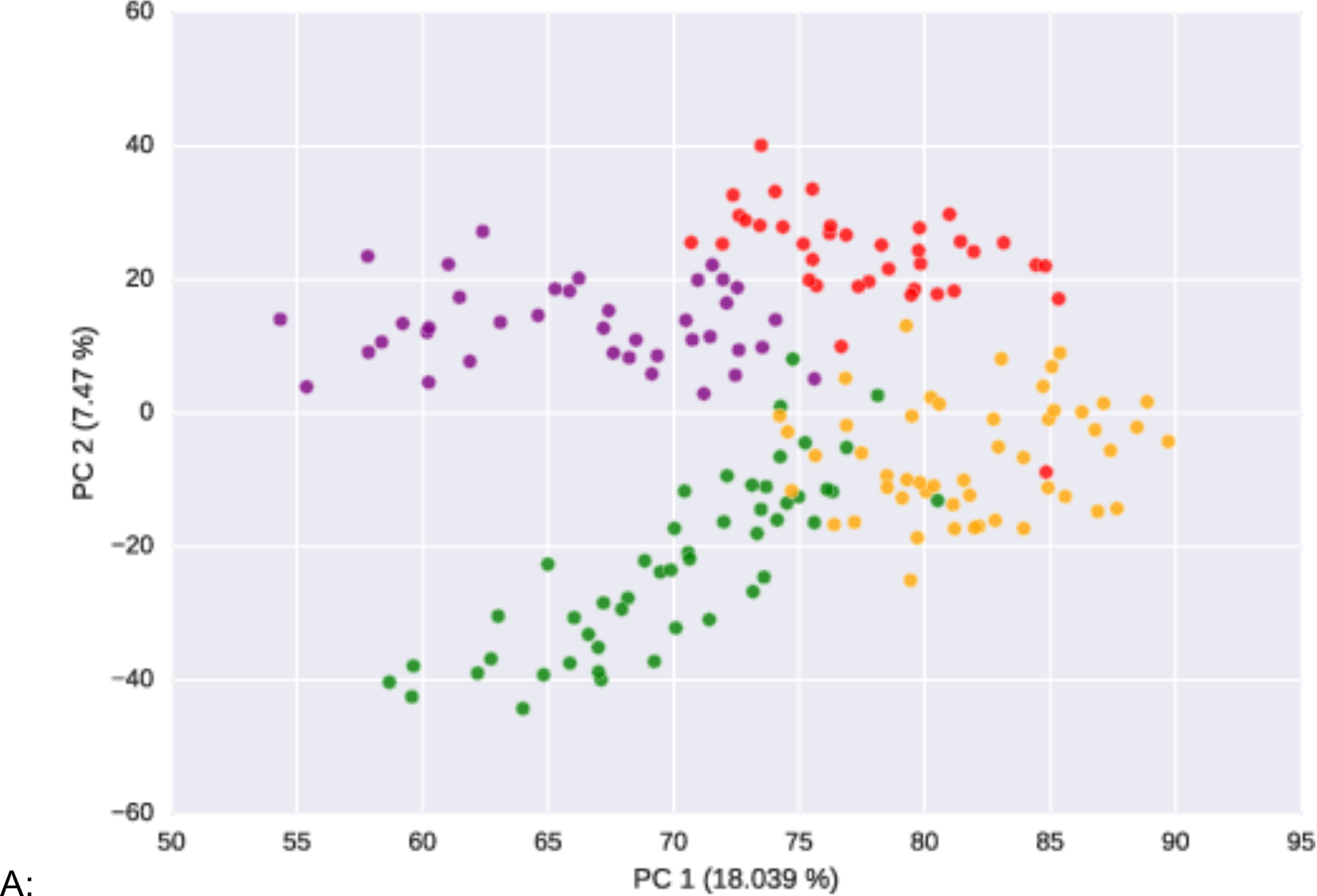
(A) Principal co-ordinate analysis of metabolome profiles inferred from 175 metagenomes from four different body-sites, displaying body-site specific metabolomes. (B-E) Trajectories of the semi-Markov chain search through the metabolome solution space towards the attractor domain of oral (B), skin (C), stool (D), and vaginal (E) metagenomes, when for each, the bottom-up ecology algorithm was initialized with two predicted metabolomes from the other body-sites.

**B:**
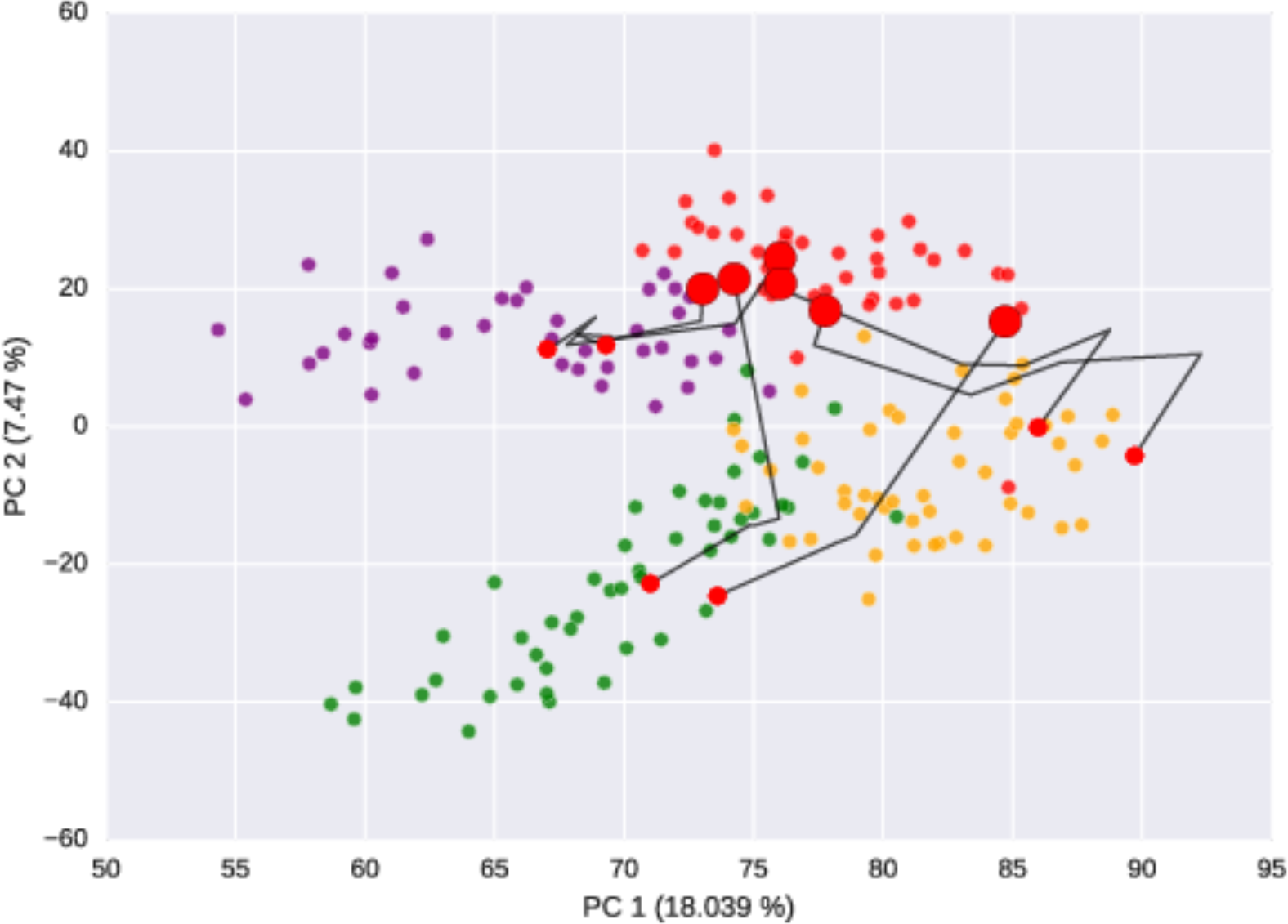

**C:**
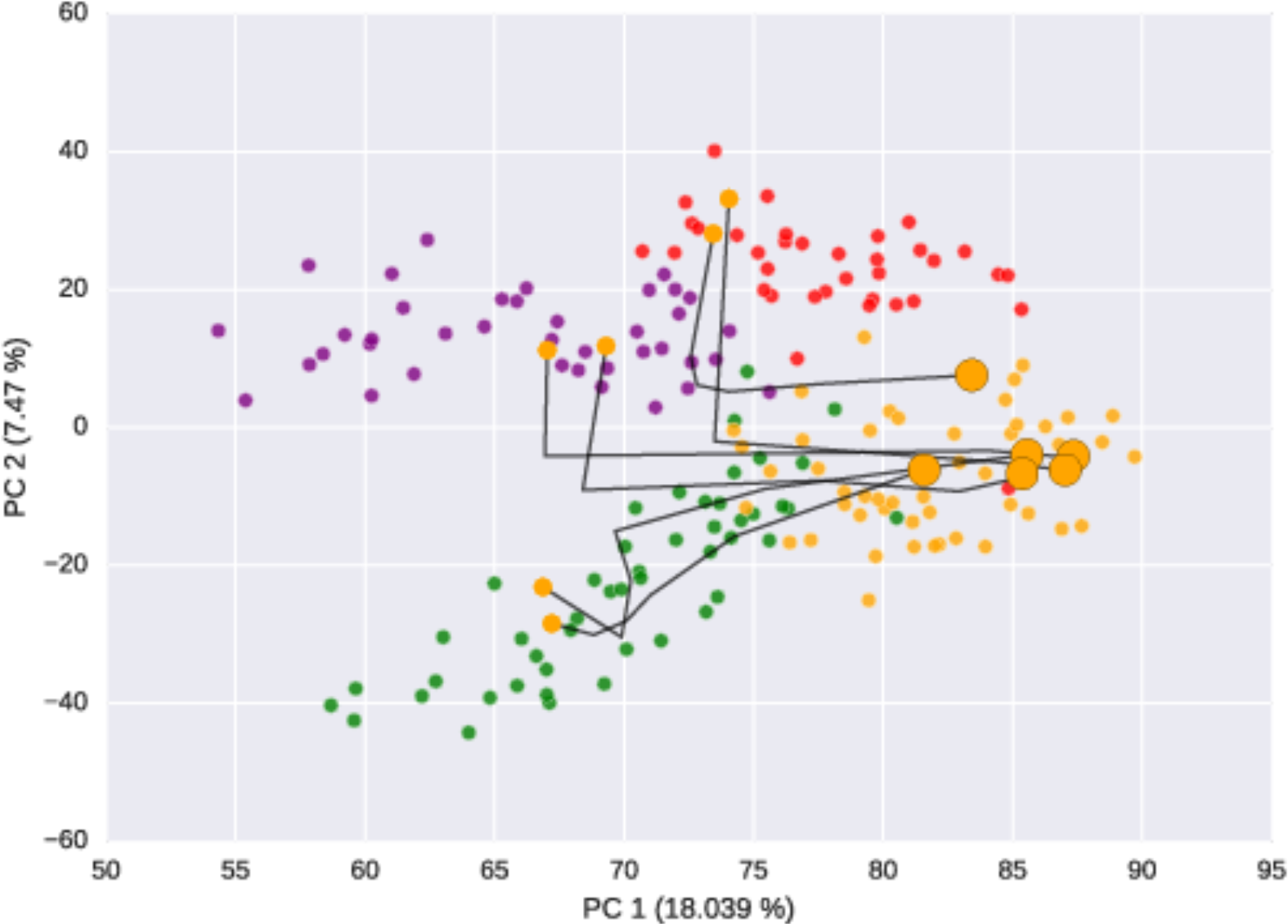

**D:**
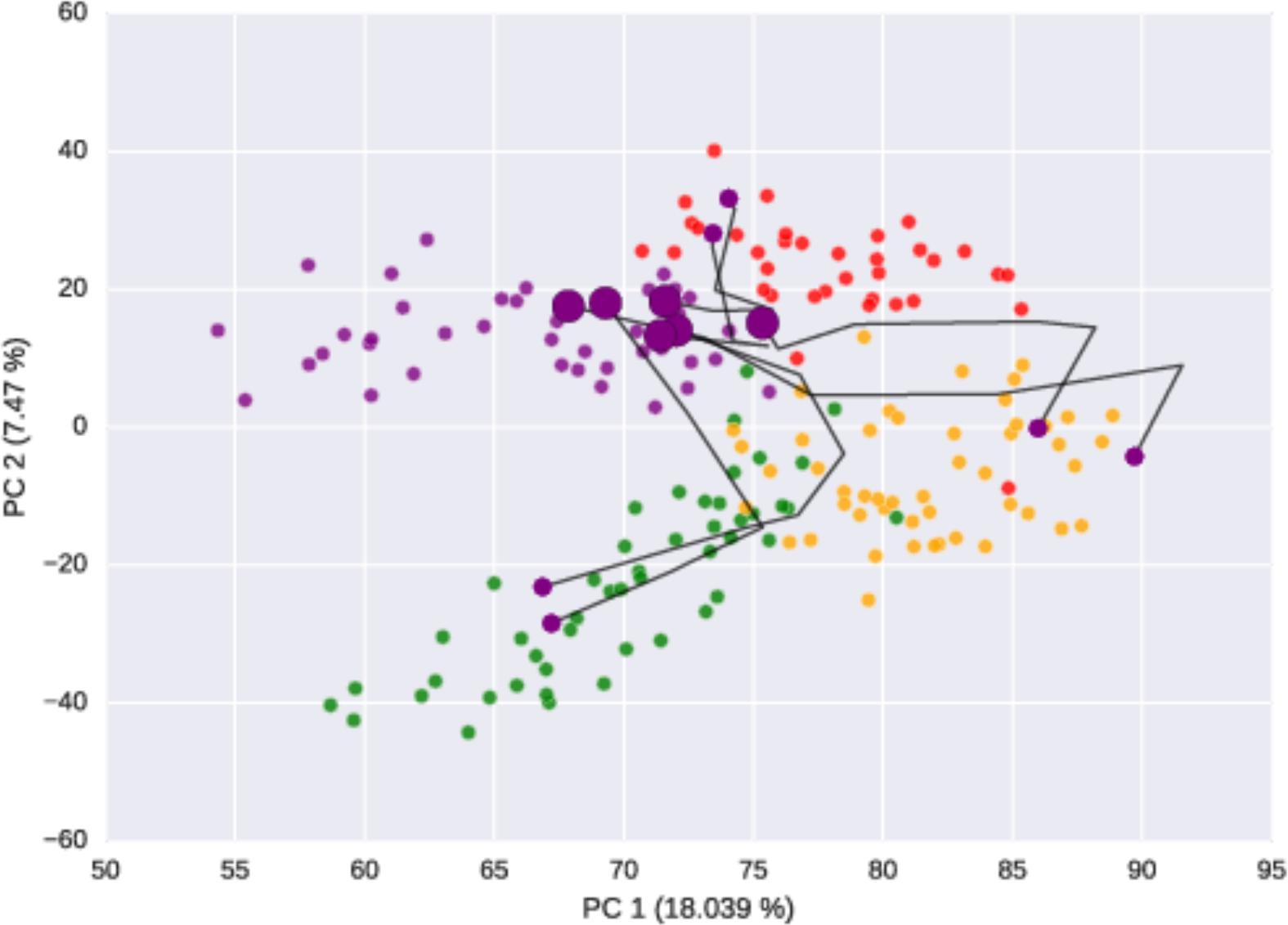

**E:**
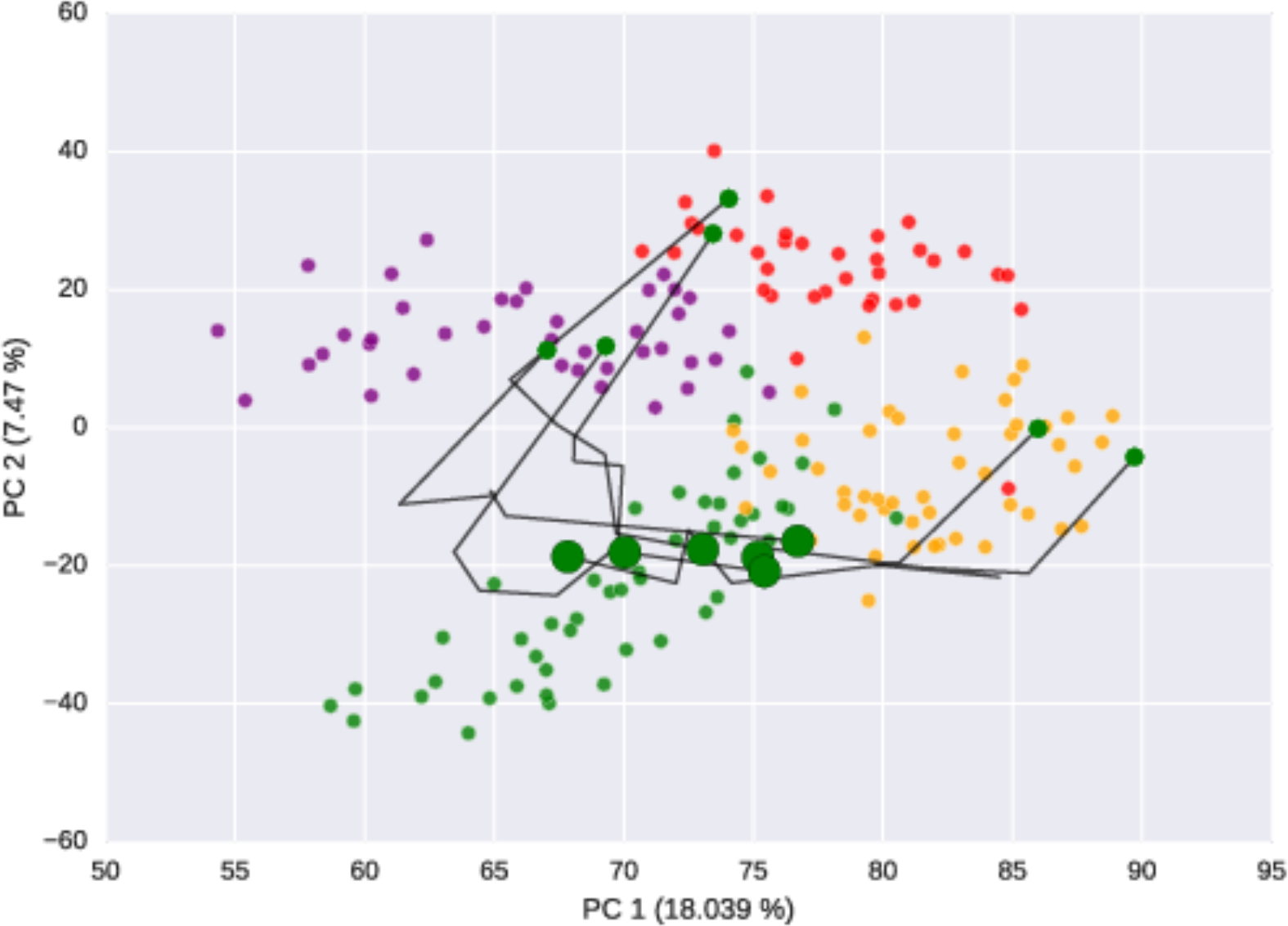

**Supplementary Figure 2.**
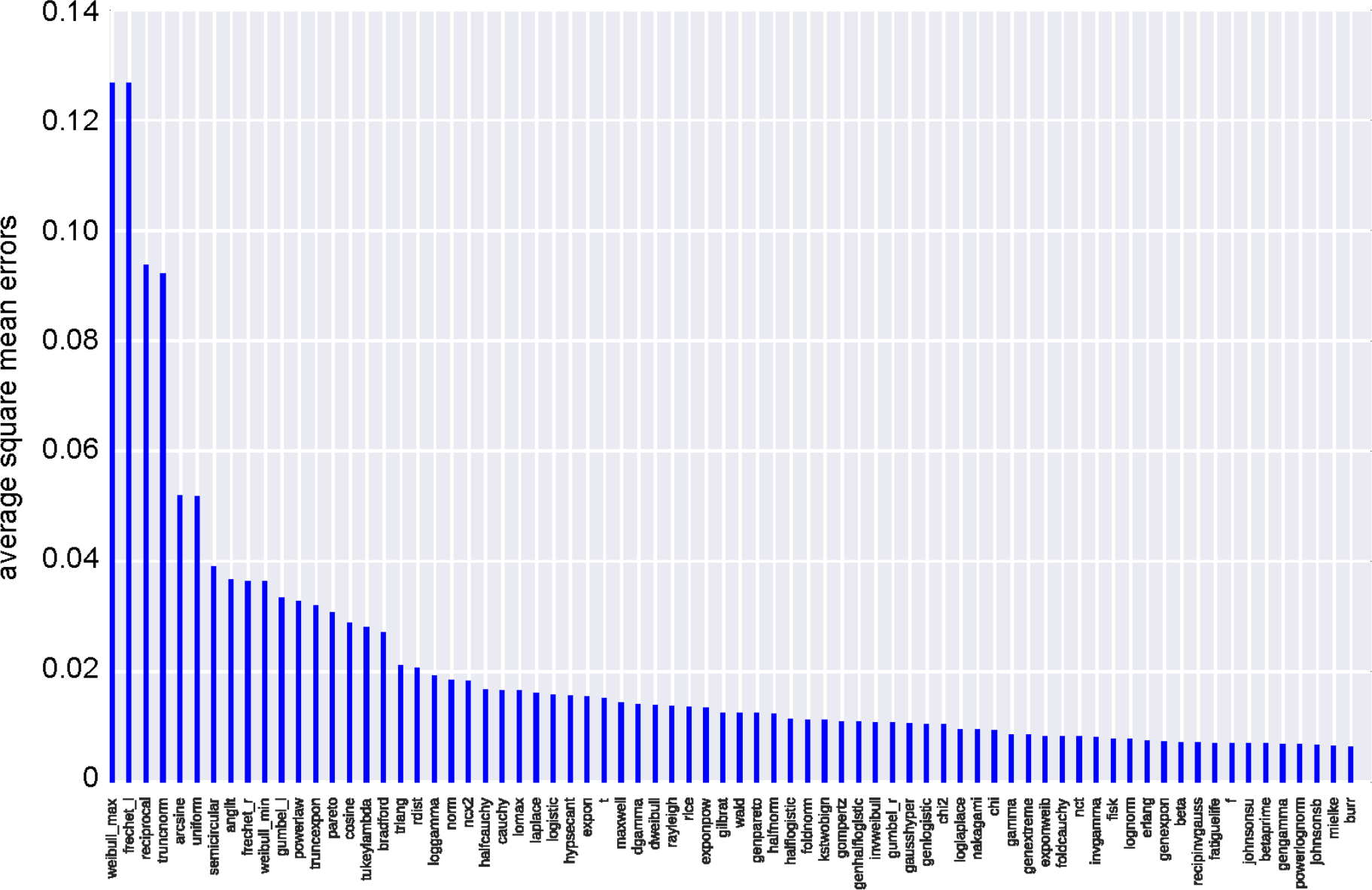
Average square mean errors of 74 common statistical distributions used to fit the distribution of metabolite abundances in all 175 predicted metabolomes.

*Supplementary File 1.* Metabolomic profiles predicted based on 37 oral, 50 skin, 39 stool, and 49 vaginal metagenomes, and six experimentally measured metabolomic profiles.

*Supplementary File 2.* Pearson correlations between 175 predicted metabolomic profiles and six measured metabolomic profiles. Correlations are only shown if >5 metabolites of the predicted metabolites were measured and *vice versa.*

**Supplementary Table 1.** Top 20 metabolites predicted by applying the bottom-up ecology algorithm to 50 human skin metagenomes. Complete lists of all predicted metabolomes for 175 metagenomes are provided in Supplementary File 1.

| Metabolite ID | Chemical | Name | Predicted relative |
| --- | --- | --- | --- |
| EX_cpd03847_e0 | C<sub>14</sub>H<sub>27</sub>O<sub>2</sub> | Myristic acid | 0.74 % |
| EX_cpd04097_e0 | Pb | Pb | 0.73 % |
| EX_cpd11590_e0 | C<sub>8</sub>H<sub>15</sub>N<sub>2</sub>O<sub>3</sub>S | met-L-ala-L | 0.72 % |
| EX_cpd00137_e0 | C<sub>6</sub>H<sub>5</sub>O<sub>7</sub> | Citrate | 0.71 % |
| EX_cpd00305_e0 | C<sub>12</sub>H<sub>17</sub>N<sub>4</sub>OS | Thiamin | 0.70 % |
| EX_cpd00276_e0 | C<sub>6</sub>H<sub>14</sub>NO<sub>5</sub> | GLUM | 0.70 % |
| EX_cpd11576_e0 | C<sub>5</sub>H<sub>11</sub>NO<sub>3</sub>S | L-methionine R-oxide | 0.70 % |
| EX_cpd00355_e0 | C<sub>11</sub>H<sub>14</sub>N<sub>2</sub>O<sub>8</sub>P | Nicotinamide ribonucleotide | 0.70 % |
| EX_cpd08305_e0 | C<sub>8</sub>H<sub>16</sub>NO | crotonobetaine | 0.70 % |
| EX_cpd03279_e0 | C<sub>10</sub>H<sub>12</sub>N<sub>4</sub>O<sub>4</sub> | Deoxyinosine | 0.70 % |
| EX_cpd00122_e0 | C<sub>8</sub>H<sub>15</sub>NO<sub>6</sub> | N-Acetyl-D-glucosamine | 0.70 % |
| EX_cpd00041_e0 | C<sub>4</sub>H<sub>6</sub>NO<sub>4</sub> | L-Aspartate | 0.70 % |
| EX_cpd00129_e0 | C<sub>5</sub>H<sub>9</sub>NO<sub>2</sub> | L-Proline | 0.69 % |
| EX_cpd00107_e0 | C<sub>6</sub>H<sub>13</sub>NO<sub>2</sub> | L-Leucine | 0.69 % |
| EX_cpd00322_e0 | C<sub>6</sub>H<sub>13</sub>NO<sub>2</sub> | L-Isoleucine | 0.69 % |
| EX_cpd01329_e0 | C<sub>36</sub>H<sub>62</sub>O<sub>31</sub> | Maltohexaose | 0.69 % |
| EX_cpd03422_e0 | C<sub>48</sub>H<sub>75</sub>CoN<sub>11</sub>O<sub>8</sub> | Cobinamide | 0.68 % |
| EX_cpd00108_e0 | C<sub>6</sub>H<sub>12</sub>O<sub>6</sub> | Galactose | 0.68 % |
| EX_cpd00084_e0 | C<sub>3</sub>H<sub>7</sub>NO<sub>2</sub>S | L-Cysteine | 0.68 % |
| EX_cpd00054_e0 | C<sub>3</sub>H<sub>7</sub>NO<sub>3</sub> | L-Serine | 0.68 % |

